# Type VI secretion system activity at lethal antibiotic concentrations leads to overestimation of weapon potency

**DOI:** 10.1101/2025.03.11.642472

**Authors:** William P. J. Smith, Elisa T. Granato

**Affiliations:** Division of Evolution, Infection and Genomics, Faculty of Biology, Medicine and Health, University of Manchester, United Kingdom; Department of Biology, University of Oxford, Oxford, United Kingdom

## Abstract

Competition assays are a mainstay of modern microbiology, offering a simple and cost-effective means to quantify microbe-microbe interactions *in vitro*. Here we demonstrate a key weakness of this method that arises when competing microbes interact via toxins, such as those secreted via the type VI secretion system (T6SS). Timelapse microscopy reveals T6SS-armed *Acinteobacter baylyi* bacteria can maintain lethal T6SS activity against *E. coli* target cells, even under selective conditions intended to eliminate *A. baylyi*. Further, this residual killing creates a density- and T6SS-dependent bias in the apparent recovery of *E. coli*, leading to a misreportin*g of* competition outcomes especially where target survival is low. We also show that incubating *A. baylyi / E. coli* co-cultures in liquid antibiotic prior to selective plating can substantially correct this bias. Our findings demonstrate the need for caution when using selective plating as part of T6SS competition assays, or assays involving other toxin-producing bacteria.

## Introduction

“Cut off a wolf’s head and it still has the power to bite”

**Hayao Miyazaki, もののけ姫 [Princess Mononoke: The First Story]**

Practiced for over a century, colony-forming unit (CFU) counting is a fundamental microbiological technique for determining the density of bacterial cells in a sample^1^. A common application for this technique is in profiling the potencies of bacterial anti-competitor toxins, as part of a killing assay^2^. Here, a “susceptible” target strain is mixed with a weapon-bearing “attacker” strain. After a set incubation period, during which the attacker kills the susceptible strain, the strain mixture is recovered, diluted, and transferred to selective media for CFU counting. The number of susceptible cells counted on media selective for that cell type is then used as a quantitative estimate of weapon potency and/or weapon susceptibility^3,4^.

A key assumption of this workflow is that killing due to toxin secretion occurs primarily during the co-culture phase, rapidly ceasing once strain mixtures are transferred to selective media. However, some anti-competitor killing mechanisms operate on fast timescales relative to the bactericidal or bacteriostatic effects of the selective media. These fast-killing mechanisms include the type VI secretion system (T6SS), a versatile anti-competitor weapon found in a broad range of gram-negative bacteria^5,6^. Resembling a poison-tipped speargun^6^, the T6SS can inject toxic effector proteins directly into neighbouring target cells, resulting in rapid (<10 min) lysis^7^.

These observations prompted us to ask: when using CFU counting in a T6SS killing assay, can interbacterial antagonism continue despite lethal selection on antibiotic-containing media? Does this influence the apparent outcome of the competition? To address these questions, we used fluorescence microscopy to measure the extent and impact of T6SS activity in *Acinetobacter baylyi* cells grown on selective media. We found that, surprisingly, T6SS activity was not detectably reduced on short (10-20 min) timescales during lethal selection. This residual activity proved sufficient to kill the majority of co-plated susceptible *Escherichia coli* cells within minutes.

We then devised a simple “ground truth” experiment, mixing T6SS attacker and susceptible strains in known ratios, and plating these on selective media to mimic the workflow of a typical killing assay. This showed that unwanted post-selection killing introduces significant biases in assay outcomes, particularly where attackers greatly outnumber susceptible cells. Biases are increased the longer cells spend in suspension before plating, but are substantially mitigated if an antibiotic pretreatment is used. Our results demonstrate that post-selection killing via contact-dependent weaponry can introduce significant biases in killing assays, leading to overestimation of weapon potency, or underestimation of resistance. Based on these findings, we recommend a cautious approach when conducting and interpreting CFU-based killing assays.

## Methods

### Bacterial strains and culture conditions

The bacterial strains used in this study are listed in Table 1. All strains were routinely cultured from frozen stocks (25% v/v glycerol) in 5 mL LB (per L MilliQ water: 10g tryptone, 10g NaCl, 5g yeast extract) supplemented with antibiotics (50 μg/mL kanamycin sulfate for *E. coli*, 50 μg/mL streptomycin sulfate for *A. baylyi* strains) in 50 mL polypropylene conical tubes. Strains were routinely pre-cultured overnight in shaking incubators (180 rpm, 16-18 h) at 37°C (*E. coli*) or at 30°C (*A. baylyi*), with lids taped ¼ turn loose to allow aeration. Optical density of liquid cultures was measured at 600 nm (OD_600_). For CFU counting, we used 120 mm vented square plates filled with ca. 40 mL molten LB agar (1.5% w/v), supplemented with antibiotics where necessary, as noted below. For time-lapse microscopy experiments, samples were kept at 30°C throughout the duration of the experiment using a custom-built incubation chamber.

**Table 1.**
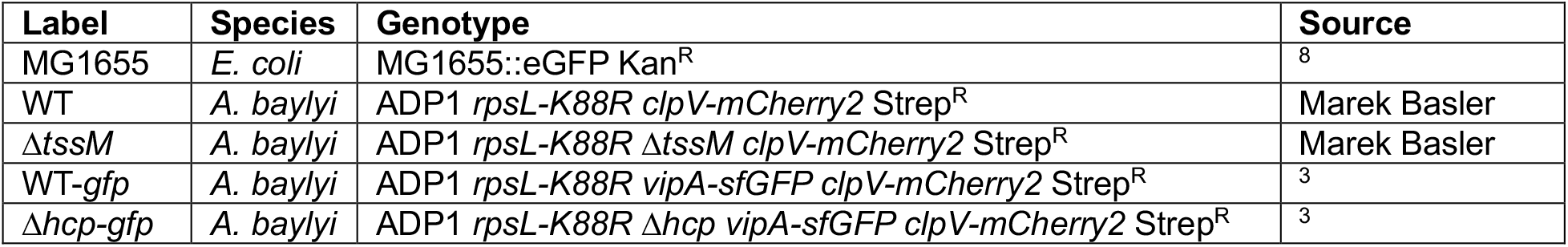
Bacterial strains used in this study.

| Label | Species | Genotype | Source |
| --- | --- | --- | --- |
| MG1655 | *E. coli* | MG1655::eGFP Kan<sup>R</sup> | <sup>8</sup> |
| WT | *A. baylyi* | ADP1 *rpsL-K88R clpV-mCherry2* Strep<sup>R</sup> | Marek Basler |
| ΔtssM | *A. baylyi* | ADP1 *rpsL-K88R* ΔtssM *clpV-mCherry2* Strep<sup>R</sup> | Marek Basler |
| WT-*gfp* | *A. baylyi* | ADP1 *rpsL-K88R vipA-sfGFP clpV-mCherry2* Strep<sup>R</sup> | <sup>3</sup> |
| Δhcp-*gfp* | *A. baylyi* | ADP1 *rpsL-K88R* Δhcp *vipA-sfGFP clpV-mCherry2* Strep<sup>R</sup> | <sup>3</sup> |

### Microscopy

#### Precultures

For images shown in Fig. 1A, overnight cultures of *A. baylyi* WT (T6SS+) were washed twice with LB, and 1 mL of washed culture was resuspended in 50 µL LB. For data and images shown in Fig. 1B-D, *A. baylyi* WT (T6SS+), Δ*tssM* (T6SS−), and *E. coli* cells were diluted into fresh LB medium from overnight cultures, grown to exponential phase (OD_600_ ∼0.6-1.0), washed twice with LB medium, and resuspended and normalized in LB medium to an OD_600_ of 1.0 (Fig. 1B) or 2.0 (Fig. 1CD).

**Fig. 1:**
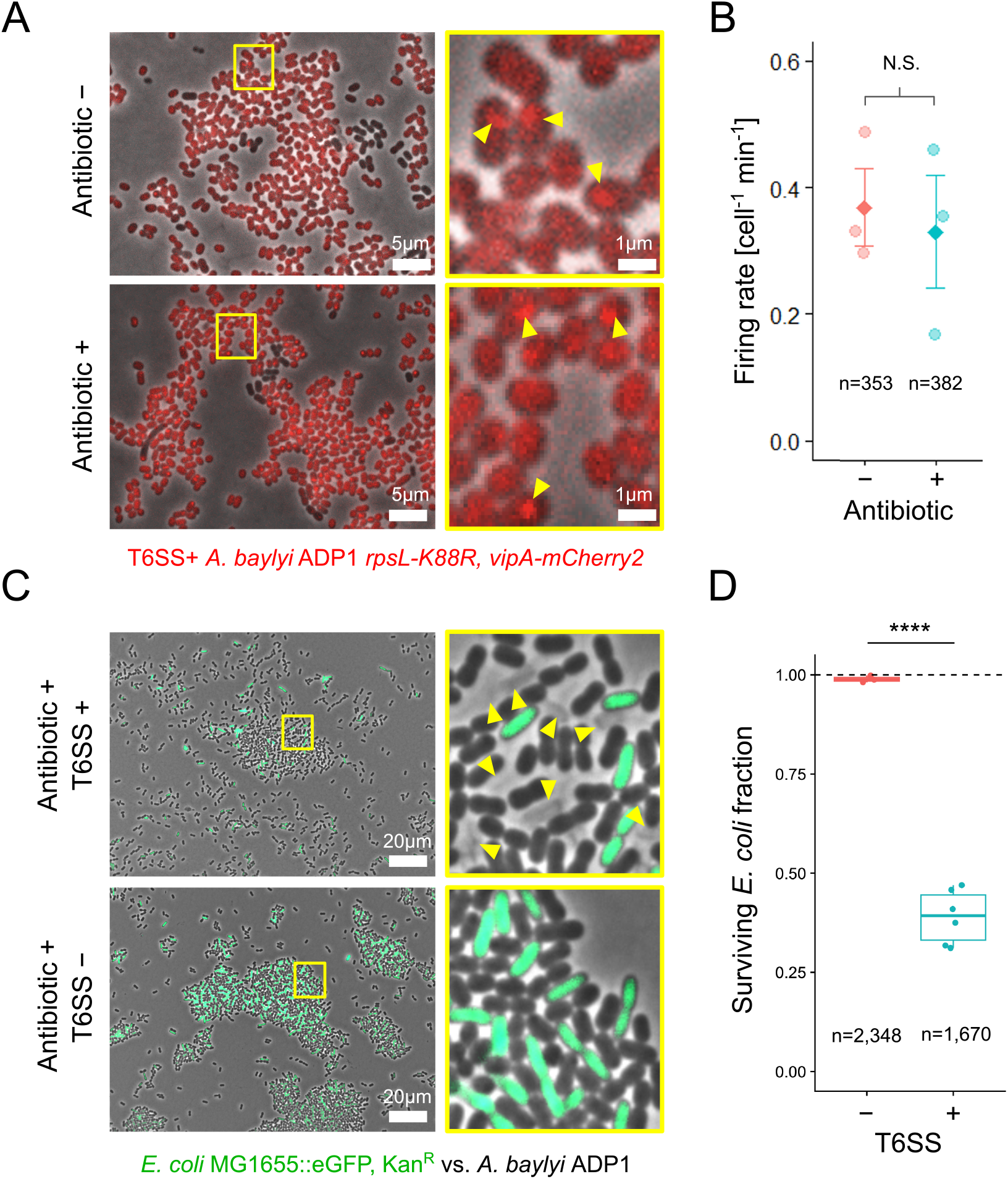
*A. baylyi* T6SS firing and *E. coli* killing are transiently maintained under lethal antibiotic selection. (**A)** Top row: T6SS+ *A. baylyi* bacteria growing on LB agarose; yellow box in left panel corresponds to zoomed section on right. Fluorescent mCherry-tagged T6SS sheaths (red; yellow arrows) are visible as dynamic foci. Bottom row: as top row, but under lethal selection (100 μg/mL kanamycin). **(B)** Quantification of T6SS firing rate (counting ClpV foci) 10-20 min after spotting on LB agarose, based on 18 images across 3 independent experiments. N.S.: non-significant (Wilcoxon rank sum test, n = 3, W = 5; p = 1.0). **(C)** Top row: *A. baylyi* T6SS+ (unlabelled) in 1:1 co-culture with GFP-labelled *E. coli* bacteria (green). Yellow box in left panel corresponds to zoomed section on right. The outlines of lysed *E. coli* cells are visible, enabling quantification of killing within a 10 min timeframe. **(D)** Measurements of surviving *E. coli* fraction (live cells / (live + dead cells)) within a 10 min timeframe for T6SS- and T6SS+ *A. baylyi* co-cultures (Welch’s two sample one-sided t-test, t = -21.26, df = 5.34, ****: p-value < 0.0001).

#### Sample preparation

*E. coli* was mixed with *A. baylyi* WT (T6SS+) at a ratio of 1:1 (20 µL each; Fig. 1B) or with either WT or Δ*tssM* (T6SS−) at a ratio of 5:1 (100 µL *A. baylyi* + 20 µL *E. coli*; Fig. 1CD). For each strain or strain mixture, 1 μL of cell suspension was then spotted onto a 1% w/v LB agarose pad, supplemented with 100 µg/mL kanamycin when appropriate. Pads were left to dry for 5 min at room temperature and then inverted onto a 5-cm diameter glass bottom Petri dish with a 3-cm diameter uncoated n°1.5 glass window (MatTek Corporation), so that the cells were sandwiched between the agarose and the glass. Samples were moved to the microscope and imaged immediately.

#### Image acquisition

Fluorescence microscopy was performed using a Zeiss Axio Observer inverted microscope with a Zeiss Plan-Apochromat 63x oil immersion objective (NA = 1.4) and ZEN Blue software (version 1.1.2.0). Image acquisition began ca. 10 min after spotting on agarose pads. Images were recorded for: 15 minutes at 10-second intervals (Fig. 1A); 5 min at 10-second intervals (Fig. 1B); or just once (Fig. 1CD). Exposure times were 30-80 ms for phase contrast, 20 ms for eGFP (ex: 488 nm | em: 507 nm), and either 100 ms (Fig. 1A) or 500 ms (Fig. 1B) for mCherry2 (ex: 589 nm | em: 610 nm).

#### Image analysis – firing rate

We measured the rate of T6SS firing in a total of N = 735 *A. baylyi* cells (Fig. 1B) across three independent experiments conducted on different days, via detection of ClpV (mCherry2) fluorescent foci^7^. Each new ClpV focus was assumed to mark one T6SS contraction event in a focal cell. For each experiment, three separate fields of view at different locations on the same agarose pad were monitored for each treatment, yielding a total of 18 time-lapse image series. Image analysis was performed using FIJI 2.1.0/1.53c^9^. For each field of view, total number of cells and ClpV foci were determined each minute for 3 min via thresholding, segmentation and object counting in the phase contrast channel, and the FIJI built-in “Find maxima” function in the mCherry2 fluorescence channel, respectively.

#### Image analysis – live vs. dead

To enumerate live/dead *E. coli* cells (Fig. 1D), we analysed 9 fields of view taken from an LB agarose pad either with kanamycin supplementation (n = 6) or without (n = 3). We divided each ∼200×160 μm field of view into 16 zones of size ∼50×40 μm. *E. coli* densities within each zone varied from 0 to ca. 150 cells. Green cells were assumed to be live *E. coli*; the outlines of lysed cells lacking eGFP signal were assumed to be dead *E. coli* (see Fig. 1C for examples). Cells within zones in which either i) focus was too poor to reliably distinguish dead cell outlines, or ii) *E. coli* eGFP fluorescence was too weak to be reliably differentiated from *A. baylyi* cells, were ignored (20/144 fields). Dead cells were almost completely absent in the T6SS− samples, and almost always found in direct contact with morphologically distinct *A. baylyi* cells in the T6SS+ samples, confirming that these dead cells are the product of T6SS-mediated killing, and not some other lethal process (e.g. antibiotic selection). Live *E. coli* cell fractions were then calculated as the ratio of live:(live+dead) *E. coli* cells. We compared measurements from T6SS+ and T6SS− treatments using Welch’s two sample one-sided t-test (built-in *t*.*test* method, R^10^ version 4.3.2 (2023-10-31)).

### Ground truth experiment

#### Preparation of cell mixtures

50 μL of *E. coli* and 500 μL of *A. baylyi* WT-*gfp* (T6SS+), Δ*hcp-gfp* (T6SS−) overnight cultures were transferred into 5 mL sterile LB (respectively, 1:100 and 1:10 dilution). These cultures were supplemented with the same antibiotics as above, and incubated for a further 2.5 h to give a final OD_600_ of approximately 1.4. Then, 1.5 mL of each exponential phase culture was washed twice with 1 mL Dulbecco’s phosphate buffered saline (dPBS; 20,000 g centrifugation for 4 min). Each washed culture was then adjusted to OD_600_ = 0.25 by diluting with dPBS, before being subjected to 5x 10-fold serial dilutions (100 μL : 900 μL) in dPBS. The *E. coli* density within each dilution was measured independently by plating in sextuplet on pre-dried, pre-warmed non-selective LB agar plates (5 μL per droplet). Once these droplets had dried, CFU plates were lidded, inverted and incubated overnight at 37°C until individual CFUs could be counted. After sampling for ground truth counts, 200 μL of washed, normalised cultures of WT-*gfp* or Δ*hcp-gfp* were individually mixed with 200 μL of each *E. coli* dilution and then vortexed for 5 s, giving 1:1, 10:1, 100:1, 1,000:1, 10,000:1 and 100,000:1 *A. baylyi*: *E. coli* mixes for each *A. baylyi* treatment. In the top row of a 96-well microtiter plate, 150 μL of each mixture was then combined with either 150 μL dPBS or 150 μL dPBS + 50 μg/mL kanamycin. Each plate was wrapped in Parafilm and incubated at room temperature (ca. 20°C).

#### CFU counting

After incubating plates for either 45 min, 1 h 45 min, 2 h 45 min, or 20 h (overnight), we counted the apparent *E. coli* and *A. baylyi* CFU density in each mixture, plating an independent dilution series for each mixture and time point (7x 10-fold serial dilutions in dPBS or dPBS + 50 μg/mL kanamycin; 5 μL of each dilution plated in triplicate on pre-dried, pre-warmed selective LB agar). To measure strain recovery, we used LB agar supplemented with 50 μg/mL kanamycin (for *E. coli*) or 100 μg/mL streptomycin (for *A. baylyi*, shown in Fig. S1). Once droplets had dried, selection plates were lidded, inverted and incubated overnight at 37°C (for *E. coli*) or 30°C (for *A. baylyi*) until individual colonies could be counted (ca. 12-18h).

#### Statistical analysis

To compare *E. coli* CFUs in ground truth, T6SS+ and T6SS− treatments, we fitted a linear mixed effects model using R’s^10^ *nlme* package (max. log likelihood fitting with Attacker strain, Pretreatment, Dilution and Dilution squared as fixed effects; plating replicate as random effect), at each experimental timepoint. We then used ANOVA to compare fixed effect sizes and their interactions and significances within each fitted model.

## Results

### *A*. *baylyi* attackers maintain T6SS firing during lethal antibiotic selection

First, we wanted to test whether *A. baylyi*, a model organism often used to study T6SS weapon function^3,7,11–14^, could still fire its T6SS apparatus under lethal selection conditions. To do this, we used single-cell microscopy to image *A. baylyi* cells labelled with a fluorescent *clpV-mCherry2* tag. The protein ClpV is an essential component of the T6SS, and ClpV foci can provide dynamic information on the rate of T6SS firing^3,7,14^. By counting ClpV foci per *A. baylyi* cell (Fig. 1A) over time, we enumerated T6SS firing events in *A. baylyi* cells grown in co-culture with *E. coli* in the absence and presence of lethal anti-*A. baylyi* antibiotic selection (0 or 100 µg/mL kanamycin). Surprisingly, we found that, ca. 10-20 min post exposure to the antibiotic, T6SS firing rate showed no detectable change compared to an antibiotic-free control (Fig. 1B; Wilcoxon rank sum test, p = 1.0). This demonstrates that, on short timescales, T6SS activity is maintained even under lethal antibiotic selection.

### Residual T6SS activity kills *E*. *coli* on selective media

Next, we wanted to test whether post-selection T6SS activity could kill a susceptible competitor strain, under antibiotic conditions that selected against the T6SS attacker. Imaging cocultures ca. 10-15 min post-mixing, we observed extensive killing of *E. coli* cells, evident from rapid lysis and concurrent loss of eGFP signal (Fig. 1C^3^). After lysis, *E. coli* cells remained visible as faint silhouettes, corresponding to the remnants of compromised cell envelopes^3^. By counting the number of intact and lysed *E. coli* cells across multiple views of the same co-culture sample, we estimated the extent of T6SS-mediated killing. Under selective conditions lethal to *A. baylyi* (100 µg/mL kanamycin), we observed that >50% of *E. coli* cells died after 10-15 min of coculture with T6SS+ *A. baylyi* attackers (Fig. 1D). By contrast, lysis rates were minimal in a T6SS− control (Fig. 1D), indicating that *E. coli* killing was indeed the product of T6SS activity. This rapid killing is consistent with previous observations of T6SS dynamics in *A. baylyi* and other species^3,7,15,16^. Crucially, it shows that significant T6SS killing is possible even under antibiotic selection conditions that are lethal to the attacker strain, highlighting the potential for induced bias in killing assay outcomes.

### Post-selection killing creates density-dependent bias in apparent competition outcomes

To test whether this effect could bias the apparent outcome of a killing assay, we created a series of *A. baylyi* / *E. coli* mixtures, with fixed *A. baylyi* density and variable *E. coli* density. Shown in Fig. 2A, these mixtures simulate the possible outputs of a typical killing assay, with susceptible cell survival varying from maximal (highest *E. coli* density, ∼2.4 × 10^7^ CFU/mL) down to the assay’s detection limit (lowest *E. coli* density, ∼200 CFU/mL). Crucially, these mixtures are of known composition, enabling comparison of measured CFU counts with known “ground truth” values. Fig. 2B compares ground-truth CFU measurements (before mixing with *A. baylyi)* with observed *E. coli* CFUs (after mixing with *A. baylyi*) for both T6SS+ and T6SS− mixtures. Following 45 min of incubation in dPBS buffer (due to the processing time of dilutions), observed *E. coli* CFUs were significantly reduced compared to ground-truth counts in mixtures with T6SS+ attackers (p = 0.02, ANOVA with linear mixed effects model, see Methods). The magnitude of this deviation depended on the *E. coli* density, with intermediate densities leading to the greatest discrepancies. We also confirmed that discrepancies disappeared in controls where T6SS was inactivated (*A. baylyi Δhcp-gfp*, p = 0.34), demonstrating that these depletion effects arise via T6SS activity. Further, increasing the incubation period to 1.5 h or 2.5 h progressively exacerbated discrepancies (Fig. 2B), suggesting that i) T6SS killing also occurs while cells are suspended in dPBS buffer, and/or ii) preincubation enhances residual killing on solid selective media. In contrast: when cell mixtures were left overnight (∼18 h) before plating, we saw substantially reduced *E. coli* survival in both T6SS+ and T6SS− mixtures, suggesting that long-term incubation in dPBS reduces *E. coli* viability independently of T6SS activity. Overall, these results show that residual T6SS activity on selective media can lead to time- and density-dependent errors in apparent killing.

**Fig. 2:**
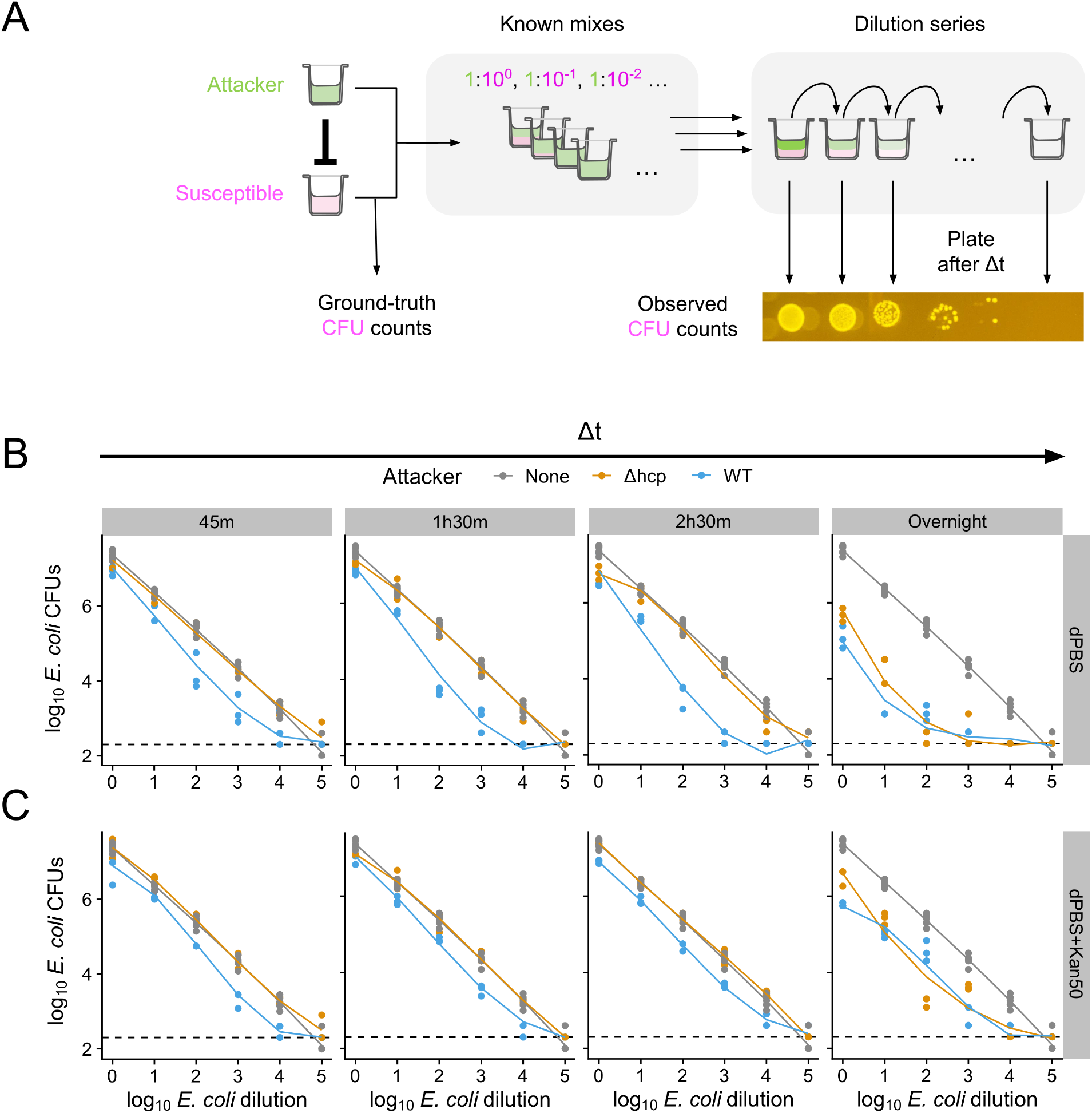
T6SS killing on selective media creates density-dependent bias in competition assays. **(A)** Schematic of ground truth experiment, showing example dilution series plated on selective media. After creation, dilution series are incubated at room temperature for variable time interval Δt, in the absence / presence of antibiotics, before plating on selective media for CFU counting. **(B)** Measurements of apparent *E. coli* recovery (CFU/mL) in co-culture with T6SS− (“Δ*hcp”*, orange) or T6SS+ (“WT”, blue) *A. baylyi* attackers, compared with known CFUs determined from monocultures (“None”, grey, replotted on each panel). Panels correspond to increasing incubation times Δt. Circles show individual datapoints; lines correspond to linear mixed effect model (fitting: maximum log likelihood, see Methods). Dashed lines show the detection limit (200 CFU/mL). (**C)** As (B), but for same mixtures incubated in dPBS + kanamycin (50 µg/mL) antibiotic pretreatment during Δt. N = 3 pseudobiological replicates (independent platings of the same mixture) per condition (N = 6 for ground truth CFU measurements).

### Preincubation with antibiotics reduces bias arising from post-selection killing

While concerning, these results also suggested a potential intervention to mitigate viability discrepancies associated with T6SS killing. Bacterial cell suspensions are typically non-conducive to T6SS killing because they do not allow for prolonged cell-cell contact^17^. We therefore reasoned that, if our *A. baylyi* + *E. coli* mixtures were incubated with anti-*A. baylyi* antibiotics before plating, this would help to remove *A. baylyi* and thus prevent the T6SS killing we identified on solid selective media. To test this hypothesis, we repeated our assay, this time incubating cell mixtures in dPBS + 50 µg/mL kanamycin, before plating on selective agar as before. While insufficient to completely remove discrepancies, this intervention was partially successful: i) in reducing that overall discrepancy between T6SS+ and T6SS− treatments (particularly for longer incubation periods), and ii) in smoothing the density dependence such that all *E. coli* densities showed a similar (minor) deviation from ground truth values. We also showed that this antibiotic pre-treatment indeed reduces *A. baylyi* viability in a time-dependent manner (Fig. S1). In sum, these results show that antibiotic exposure prior to selective plating can mitigate viability discrepancies.

## Discussion

Co-culture assays are frequently used to analyse microbial interactions^3,4,7,18–27^. The outcome of these assays is often evaluated using selective plating and CFU counting, allowing co-cultured strains to be distinguished and enumerated based on their selective markers^28^. These techniques offer a simple and cost-effective way to quantify a focal strain’s growth or survival in the presence of other microorganisms. CFU counting also affords a very high dynamic range (typically >8 orders of magnitude), enabling viable cell counts from samples with very high or very low cell densities^29^. In studies aimed at estimating the anti-competitor effects of microbial weapons, such as T6SSs, CFU counting has accordingly been the gold-standard evaluation method for decades^28,30^.

Here, we show that T6SS-mediated killing continues even when cell mixtures are plated on selective media intended to disable attackers (Fig. 1). We also demonstrate that this effect leads to a significant and density-dependent bias in the apparent survival of the targeted strain (Fig. 2, Fig. S1). Importantly, the presence of residual killing on selective media does more than simply extend the effective competition period. This is because of the coupling between susceptible survival and attacker density in the CFU counting dilution series: the fewer the surviving bacteria, the lower the dilution at which they are detected, and so the higher the attacker density present within the corresponding CFU spot.

Residual killing on selective media thus has the potential to introduce large (>10-fold) errors to in competition outcome measurements. Specifically, the effects of high-potency weapons could be *overestimated*, since i) competition is extended beyond the intended interaction time, and ii) transferring cells to selective media generally involves resuspension and re-assortment, further increasing killing^31^. For the same reasons, *resistance* to weapon attacks could be *underestimated*, since residual killing can significantly reduce apparent survival of the target strain, post-competition^4,8,26^.

Our findings have several limitations. First, while *A. baylyi* / *E. coli* cocultures are an established model for understanding T6SS interactions^3,7,11–14^, this is but one example of a lethal microbe-microbe interaction. Other attacker-target pairings may introduce additional factors that modulate intermicrobial competition (e.g. differential adhesion^21,32^, density-dependent toxin production^33,34^, potentially introducing other biases alongside those shown here. Our findings are also limited to a specific type of selection (kanamycin); other selective media may result in different residual killing profiles. We speculate that bacteriostatic, colorimetric or non-auxotrophic media may be particularly vulnerable to residual killing biases, since these would be expected to disable attacker strains less rapidly (or not at all) compared with media containing bacteriolytic antibiotics. Conversely, it is possible that some CFU counting methods may be intrinsically less prone to bias than others. The recently-published geometric volumetric analysis (GVA) method^29^ counts CFUs by embedding cells in agarose, rather than growing them on top of it, and so may mitigate biases by physically separating attacker and target cells. Further work is required to test these possibilities.

Immediately however, our findings emphasise the need for caution when interpreting the results of co-culture CFU counts, especially when lethal microbe-microbe interactions are (or may be) involved. We speculate that our findings could extrapolate to other rapid killing mechanisms, including other contact weapons^32–34^ and diffusible toxins^35,36^. Based on our findings we recommend that, where co-cultured microbes are known to interact (particularly via potent antimicrobials), researchers perform ground-truth tests to check for biases. Where biases are present, we showed that these may be partially alleviated by incubating cell suspensions with antibiotics before plating. Failing this, alternative measures of competition outcome where cells are physically separated (e.g. flow cytometers^37^, qPCR^38^) or enumerated *in situ* (barcode sequencing^39^, quantitative fluorescence microscopy^14,40^ or non-selective colorimetric killing assays^3^ may become advantageous.

CFU counting has many strengths and is likely to remain a common technique for quantifying the outcomes of competition assays. However, our work warns against uncritical use of this method, as residual killing effects can lead to misleading results where competition involves fast-killing antimicrobials.

## Acknowledgements

We thank Marek Basler and Mike Brockhurst for providing the strains used in this study. We are grateful to Chris Knight for his advice on the use of the *nlme* package, and to Katie MacGillivray and Sean Booth for providing feedback on our manuscript draft.

