## Supplementary Information for "Type VI secretion system activity at lethal antibiotic concentrations leads to overestimation of weapon potency"

### Supplementary Figures

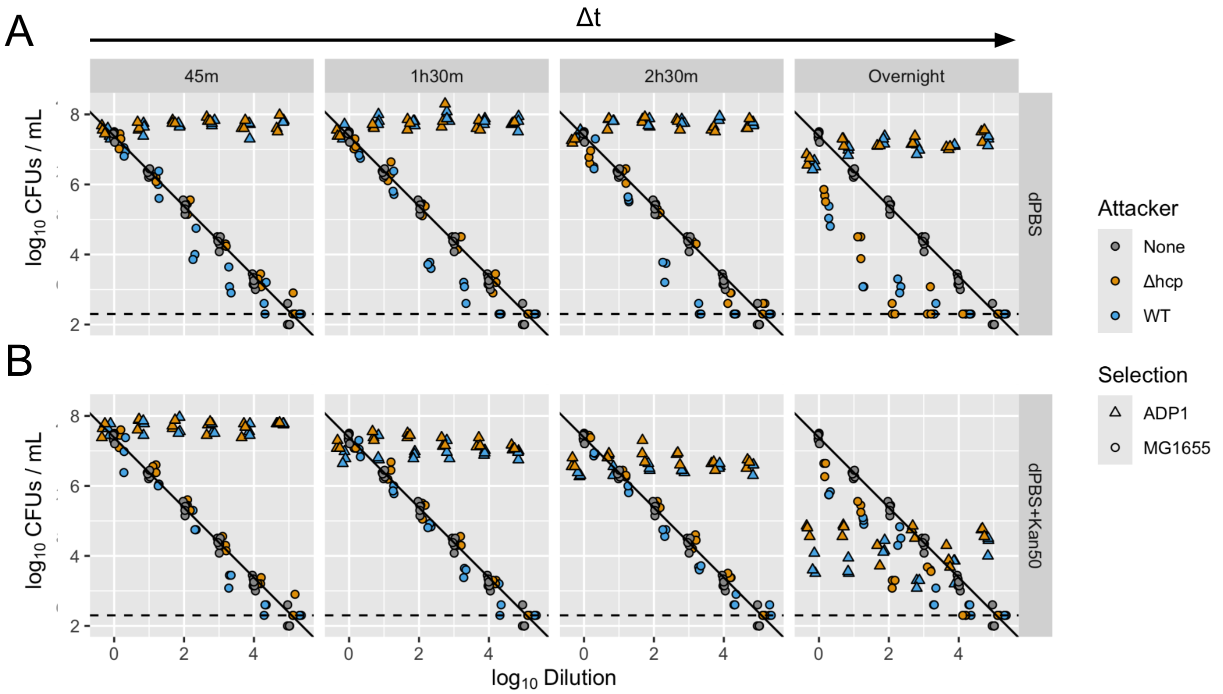


**Fig. S1: Raw data for "Ground truth" experiment shown in Fig 1. (A)** Measurements of apparent *E. coli* ("MG1655", circles) and *A. baylyi* ("ADP1", triangles) recovery are coloured according to attacker type ("Δhcp": T6SS−; "WT", T6SS+), and plotted alongside known *E. coli* CFUs determined from monocultures ("None", grey, replotted on each panel). Panels correspond to increasing incubation times Δt. Dashed lines show the detection limit (200 CFU/mL); solid line shows the predicted *E. coli* recovery (CFU = Dilution) based on extrapolation from the undiluted *E. coli* sample (2.4 x $10^7$ CFU/mL). **(B)** As (A) but for same mixtures incubated in dPBS + kanamycin (50 μg/mL) antibiotic pretreatment during Δt. N = 3 pseudobiological replicates (independent platings of the same mixture) per condition (N = 6 for ground truth CFU measurements).
